## Supplementary Material for "Identification of potential inflammation markers for outgrowth of cow’s milk allergy"

[#] Shared last authorship

* Corresponding author at: Laboratory of Microbiology, Wageningen University, Wageningen, The Netherlands.

### SUPPLEMENTARY METHODS & RESULTS

#### 1. Demonstration of IgE-mediated CMA

In short, infants sensitized to CM, confirmed by CM-specific serum IgE >0.1 kU/L and/or CM skin prick test wheal size ≥3 mm, were diagnosed with CMA as follows. For some infants, CMA diagnosis was confirmed by an open or double-blind placebo-controlled CM challenge (DBPCFC). For others, diagnosis was confirmed by a history of anaphylaxis reaction to CM reported by two physicians.

#### 2. Demonstration of resolution of CMA

Infants received a DBPCFC with CM powder, and in case this was negative an oral fresh milk challenge was performed. Infants negative for both tests were considered to have outgrown their CMA.

#### 3. The proximity extension assay (PEA) protocol

In PEA, matched antibody pairs are labelled with DNA oligonucleotides. These antibody pairs bind their target proteins in the samples, which brings the DNA nucleotides in proximity to each other. The DNA nucleotides which are in proximity hybridize, and are extended by addition of a DNA polymerase. This results in the creation of DNA barcodes, which are subsequently amplified and quantified by qPCR.

### 4. NPX values and between-plate normalization

NPX is an arbitrary unit on log2 scale used by Olink®. Between-plate normalization was performed using intensity normalization v.2, which applies the median of the samples as normalization factor. More detail on this normalization method is provided on the Olink® website (https://7074596.fs1.hubspotusercontent-na1.net/hubfs/7074596/05-white%20paper%20for%20website/1096-olink-data-normalization-white-paper.pdf ).

### 5. Quality control

Quality control (QC) was performed by adding four internal controls to each sample. Individual samples pass the QC when their deviation from the median value of the controls < 0.3 NPX from the median.

### 6. Results of the quality control and filtering

Four samples (two from visit 6 months, two from 12 months) did not pass the QC. As these samples were not detected as outliers by principal component analysis (Figure S1), they were not removed from the data set. Thirty-four proteins had a value below LOD in more than 20% of the samples and were filtered out.

### 7. Linear mixed models (LMM) analysis

To determine differences between allergy groups (resolution of CMA at 12 months (yes/no)) within visits, and between visits within allergy groups, an LMM with allergy status, visit and the interaction of allergy status and visit as fixed effects and subject as random effect was fit for each protein. A post-hoc test was performed to enable pairwise comparisons. A similar mixed model analysis was performed to determine differences between treatment groups (AAF/AAF-syn) within visits, and between visits within treatment groups. The fixed effects were treatment, visit and the interaction of treatment and visit, and the random effect was subject. In all cases, p-values were adjusted for multiple hypothesis testing using the Benjamini-Hochberg procedure[1], and results with adjusted p-value ≤ 0.05 were considered significant.

### 8. Pathway analysis with ConsensusPathDB

For each protein, the mean measurement value of the samples for each of the two groups in the comparison was supplied as input. Subsequently, a paired Wilcoxon signed-rank test was carried out on each Kyoto Encyclopedia of Genes and Genomes (KEGG) pathway that included a minimum of 4 of the measured proteins. A p-value per KEGG pathway was assigned, and the p-values were corrected for multiple testing using the false discovery rate (FDR) method. The adjusted p-values are presented as q-values in the results tables, results with q-value ≤ 0.05 were considered significant.

**Table S1.** Baseline characteristics per allergy group. Numeric variables are presented as mean ± standard deviation, categorical variables are presented as number. P-values were determined by a two-sided MannWhitney U-test for numeric variables and a Fisher's exact test for categorical variables. When the p-value was already calculated previously, this is indicated in the column "reference". Significant differences between allergy groups are indicated in bold. Abbreviations: CM: cow's milk, SPT: skin prick test, AD: atopic dermatitis, SCORAD: SCORing Atopic Dermatitis.

| | **Persistent CMA at 12M (n = 15)** | **Resolved CMA at 12M (n = 24)** | **p-value** | **reference** |
|---|---|---|---|---|
| age (months) | 9.68 ± 2.63 | 8.56 ± 30.04 | 0.254 | [2] |
| sex | | | 0.477 | [2] |
| female | 3 | 8 | | |
| male | 12 | 16 | | |
| race | | | 0.617 | this study |
| Caucasian/white | 4 | 5 | | |
| Asian | 10 | 18 | | |
| other | 1 | 1 | | |
| CM-specific IgE (kU/L) | 23.28 ± 50.99 | 1.70 ± 2.80 | **< 0.001** | this study |
| CM SPT wheal size (mm) | 6.80 ± 2.65 | 4.17 ± 2.20 | **0.003** | this study |
| AD severity (SCORAD) | 16.27 ± 13.24 | 8.98 ± 14.41 | **0.036** | [2] |
| other food allergy | | | 0.333 | this study |
| yes | 9 | 10 | | |
| no | 6 | 14 | | |
| treatment | | | 1.000 | [2] |
| AAF | 6 | 10 | | |
| AAF-syn | 9 | 14 | | |

**Table S2.** Mean ± standard deviation for IgE-specific α-lactalbumin, β-lactoglobulin and casein for all infants, infants with persistent CMA and infants with resolved CMA. P-value obtained by comparing the two allergy groups with two sided Mann-Whitney U-test. Significant results are indicated in bold.

| **parameter** | **Total (n = 39)** | **Persistent CMA at visit 12M (n=15)** | **Resolved CMA at visit 12M (n=24)** | **p-value (two-sided Mann-Whitney U-test)** |
|---|---|---|---|---|
| Alpha-lactalbumin-specific IgE (kU/L) | | | | |
| baseline | 1.33 ± 2.62 | 2.11 ± 2.76 | 0.84 ± 2.46 | **0.01672** |
| 12 months | 1.06 ± 2.02 | 2.25 ± 2.81 | 0.31 ± 0.67 | **0.00113** |
| Beta-lactoglobulin-specific IgE (kU/L) | | | | |
| baseline | 2.36 ± 5.06 | 5.15 ± 7.36 | 0.62 ± 1.03 | **0.00693** |
| 12 months | 2.07 ± 4.99 | 4.98 ± 7.26 | 0.26 ± 0.40 | **0.00085** |
| Casein-specific IgE (kU/L) | | | | |
| baseline | 12.28 ± 56,85 | 30.66 ± 90.45 | 0.80 ± 1.44 | **0.00231** |
| 12 months | 6.66 ± 17.74 | 16.74 ± 26.01 | 0.35 ± 0.66 | **0.00012** |

**Ethical approval**

This multicenter study was performed according to the World Medical Association (WMA) Declaration of Helsinki and the International Conference on Harmonization guidelines for Good Clinical Practice[3]. Table S3 provides the list of all national ethics committees, institutional review boards and regulatory authorities that approved the study protocol and amendments. Written informed consent for the collection and analysis of the data was obtained from the parents of all infants included in this study[1].

**Table S3.** List of ethics committees, institutional review boards and regulatory authorities that approved this study.

| **Country** | **Ethics committees, institutional review boards and regulatory authorities** |
|---|---|
| United Kingdom | NRES Committee North East - Sunderland (Central Ethics Committee MREC) and the local R&Ds from the following hospitals: Great Northern Children's Hospital / Newcastle General Hospital; Southampton General Hospital; Guys & St Thomas; Barts / Royal London Hospital; Leicester Royal Infirmary |
| Germany | Ethikkommission Charité – Ethikausschuss 2 am Campus Virchow Klinikum; Ethikkommission Ärztekammer Nordrhein Düsseldorf, Ethik-Kommission der Medizinischen Fakultät der Ruhr Universität Bochum |
| Italy | Comitato Etico per la Sperimentazione Clinica della Province della Pronvincia di Padova; Comitato Etico per la Sperimentazione Clinica della Province di Verona e Rovigo |
| Singapore | Singhealth Centralised Institutional Review Board (CIRB) E; National Healthcare Group (NHG) Domain Specific Review Board |
| Thailand | Institutional Review Board of the Faculty of Medicine, Chulalongkorn University; Committee on Human Rights Related to Research Involving Human Subjects, Faculty of Medicine Ramathibodi Hospital, Mahidol University; Ethics Committee of the Faculty of Medicine, Prince of Songkla University |
| United States of America | Institutional Review Board of the Mount Sinai School of Medicine; Institutional Review Board for Human Subject Research for Baylor College of Medicine and Affiliated Hospitals (BCM IRB); University of Arkansas for Medical Sciences (UAMS) Institutional Review Board |

**Table S4.** List of proteins after filtering out proteins that had a value below the limit of detection in more than 20% of the samples.

| Uniprot ID | Olink ID | Symbol | Protein name |
|---|---|---|---|
| Q13541 | OID00536 | 4E-BP1 | Eukaryotic translation initiation factor 4E-binding protein 1 |
| P00813 | OID00560 | ADA | Adenosine Deaminase |
| Q14790 | OID00550 | CASP-8 | Caspase-8 |
| Q99731 | OID00513 | CCL19 | C-C motif chemokine 19 |
| P78556 | OID00556 | CCL20 | C-C motif chemokine 20 |
| P55773 | OID00530 | CCL23 | C-C motif chemokine 23 |
| Q9NRJ3 | OID00539 | CCL28 | C-C motif chemokine 28 |
| P10147 | OID00532 | CCL3 | C-C motif chemokine 3 |
| P13236 | OID00498 | CCL4 | C-C motif chemokine 4 |
| P25942 | OID00542 | CD40 | CD40L receptor |
| P06127 | OID00531 | CD5 | T-cell surface glycoprotein CD5 |
| Q9H5V8 | OID00476 | CDCP1 | CUB domain-containing protein 1 |
| P09603 | OID00562 | CSF-1 | Macrophage colony-stimulating factor 1 |
| P28325 | OID00491 | CST5 | Cystatin D |
| P78423 | OID00552 | CX3CL1 | Fractalkine |
| P09341 | OID00496 | CXCL1 | C-X-C motif chemokine 1 |
| P02778 | OID00535 | CXCL10 | C-X-C motif chemokine 10 |
| O14625 | OID00486 | CXCL11 | C-X-C motif chemokine 11 |
| P42830 | OID00520 | CXCL5 | C-X-C motif chemokine 5 |
| P80162 | OID00534 | CXCL6 | C-X-C motif chemokine 6 |
| Q07325 | OID00490 | CXCL9 | C-X-C motif chemokine 9 |
| Q8NFT8 | OID01213 | DNER | Delta and Notch-like epidermal growth factor-related receptor |
| P80511 | OID00541 | EN-RAGE | Protein S100-A12 |
| O95750 | OID00545 | FGF-19 | Fibroblast growth factor 19 |
| P49771 | OID00533 | Flt3L | Fms-related tyrosine kinase 3 ligand |
| P14210 | OID00522 | HGF | Hepatocyte growth factor |
| P01583 | OID00493 | IL-1 alpha | Interleukin-1 alpha |
| Q08334 | OID00515 | IL-10RB | Interleukin-10 receptor subunit beta |
| P29460 | OID00523 | IL-12B | Interleukin-12 subunit beta |
| Q13261 | OID00514 | IL-15RA | Interleukin-15 receptor subunit alpha |
| Q14116 | OID00501 | IL-18 | Interleukin-18 |
| Q13478 | OID00517 | IL-18R1 | Interleukin-18 receptor 1 |
| Q9UHF4 | OID00489 | IL-20RA | Interleukin-20 receptor subunit alpha |
| P05231 | OID00482 | IL-6 | Interleukin-6 |
| P13232 | OID00478 | IL-7 | Interleukin-7 |
| P10145 | OID00471 | IL-8 | Interleukin-8 |
| P01137 | OID00480 | LAP TGF-beta-1 | Latency-associated peptide transforming growth factor beta-1 |
| P15018 | OID00547 | LIF | Leukemia inhibitory factor |
| P42702 | OID00511 | LIF-R | Leukemia inhibitory factor receptor |
| P13500 | OID00484 | MCP-1 | Monocyte chemotactic protein 1 |
| P80075 | OID00549 | MCP-2 | Monocyte chemotactic protein 2 |

**Table S4.** (continued)

| Uniprot ID | Olink ID | Symbol | Protein name |
|---|---|---|---|
| Q99616 | OID00504 | MCP-4 | Monocyte chemotactic protein 4 |
| P03956 | OID00510 | MMP-1 | Matrix metalloproteinase-1 |
| P09238 | OID00527 | MMP-10 | Matrix metalloproteinase-10 |
| O00300 | OID00479 | OPG | Osteoprotegerin |
| P13725 | OID00494 | OSM | Oncostatin-M |
| Q9NZQ7 | OID00518 | PD-L1 | Programmed cell death 1 ligand 1 |
| P21583 | OID00500 | SCF | Stem cell factor |
| Q8IXJ6 | OID00538 | SIRT2 | SIR2-like protein 2 |
| O95630 | OID00558 | STAMBP | STAM-binding protein |
| P01135 | OID00503 | TGF-alpha | Transforming growth factor alpha |
| P01375 | OID05548 | TNF | Tumor necrosis factor |
| Q07011 | OID00553 | TNFRSF9 | Tumor necrosis factor receptor superfamily member 9 |
| O43557 | OID00506 | TNFSF14 | Tumor necrosis factor ligand superfamily member 14 |
| P50591 | OID00488 | TRAIL | TNF-related apoptosis-inducing ligand |
| O43508 | OID00555 | TWEAK | Tumor necrosis factor (Ligand) superfamily, member 12 |
| P00749 | OID00481 | uPA | Urokinase-type plasminogen activator |
| P15692 | OID00472 | VEGFA | Vascular endothelial growth factor A |

**Table S5.** Results of comparing visits within allergy groups using linear mixed models and post hoc analysis. Allergic = persistent CMA at visit 12M; tolerant = resolved CMA at visit 12M. Number of samples: persistent CMA – 0M: 15, persistent CMA – 6M: 15, persistent CMA – 12M: 14, resolved CMA – OM: 24, resolved CMA – 6M: 24, resolved CMA – 12M: 24.

See Excel file Supplementary_table_S5.xlsx

**Table S6.** Results of comparing allergy groups within visits using linear mixed models and post hoc analysis. Allergic = persistent CMA at visit 12M; tolerant = resolved CMA at visit 12M. Number of samples: persistent CMA – 0M: 15, persistent CMA – 6M: 15, persistent CMA – 12M: 14, resolved CMA – OM: 24, resolved CMA – 6M: 24, resolved CMA – 12M: 24.

See Excel file Supplementary_table_S6.xlsx

**Table S7.** Results of comparing visits within treatment groups using linear mixed models and post hoc analysis. AAF: standard amino acid formula; AAF-syn: amino acid formula with synbiotic blend. Number of samples: AAF – OM: 16, AAF – 6M: 16, AAF-12M: 16, AAF-syn – 0M: 23, AAF-syn – 6M: 23, AAF-syn – 12M: 22.

See Excel file Supplementary_table_S7.xlsx

**Table S8.** Results of comparing treatment groups within visits using linear mixed models and post hoc analysis. AAF: standard amino acid formula; AAF-syn: amino acid formula with synbiotic blend. Number of samples: AAF – OM: 16, AAF – 6M: 16, AAF-12M: 16, AAF-syn – 0M: 23, AAF-syn – 6M: 23, AAF-syn – 12M: 22.

See Excel file Supplementary_table_S8.xlsx

**Table S9.** Results of comparing visits within allergy groups using KEGG pathway enrichment in ConsensusPathDB. Pathways with at least 4 measured proteins are displayed. Allergic = persistent CMA at visit 12M; tolerant = resolved CMA at visit 12M. Number of samples: persistent CMA – 0M: 15, persistent CMA – 6M: 15, persistent CMA – 12M: 14, resolved CMA – OM: 24, resolved CMA – 6M: 24, resolved CMA – 12M: 24.

See Excel file Supplementary_table_S9.xlsx

**Table S10.** Results of comparing allergy groups within visits using KEGG pathway enrichment in ConsensusPathDB. Pathways with at least 4 measured proteins are displayed. Allergic = persistent CMA at visit 12M; tolerant = resolved CMA at visit 12M. Number of samples: persistent CMA – 0M: 15, persistent CMA – 6M: 15, persistent CMA – 12M: 14, resolved CMA – OM: 24, resolved CMA – 6M: 24, resolved CMA – 12M: 24.

See Excel file Supplementary_table_S10.xlsx

**Table S11.** Results of comparing visits within treatment groups using KEGG pathway enrichment in ConsensusPathDB. Pathways with at least 4 measured proteins are displayed. AAF: standard amino acid formula; AAF-syn: amino acid formula with synbiotic blend. Number of samples: AAF – OM: 16, AAF – 6M: 16, AAF-12M: 16, AAF-syn – 0M: 23, AAF-syn – 6M: 23, AAF-syn – 12M: 22.

See Excel file Supplementary_table_S11.xlsx

**Table S12.** Results of comparing treatment groups within visits using KEGG pathway enrichment in ConsensusPathDB. Pathways with at least 4 measured proteins are displayed. AAF: standard amino acid formula; AAF-syn: amino acid formula with synbiotic blend. Number of samples: AAF – OM: 16, AAF – 6M: 16, AAF-12M: 16, AAF-syn – 0M: 23, AAF-syn – 6M: 23, AAF-syn – 12M: 22.

See Excel file Supplementary_table_S12.xlsx

**Table S13.** Comparison of the number of infections (num inf). (A) Between 6-month visit intervals within allergy groups. P-values were determined by a paired two-sided MannWhitney U-test. (B) Between allergy groups within 6-month visit intervals. P-values were determined by a two-sided MannWhitney U-test. (C) Between 6-month visit intervals within treatment groups. P-values were determined by a paired two-sided MannWhitney U-test. (D) Between treatment groups within 6-month visit intervals. P-values were determined by a two-sided MannWhitney U-test.

| | comparison | | results | | |
|---|---|---|---|---|---|
| | 1 | 2 | mean1 ± sd1 | mean2 ± sd2 | p-value |
| (A) | persistent CMA at 12M num inf 0 to 6M | persistent CMA at 12M num inf 6 to 12M | 1.67 ± 1.45 | 1.33 ± 1.23 | 0.440 |
| | resolved CMA at 12M num inf 0 to 6M | resolved CMA at 12M num inf 6 to 12M | 1.25 ± 4.61 | 0.71 ± 0.81 | 0.079 |
| (B) | persistent CMA at 12M num inf 0 to 6M | resolved CMA at 12M num inf 0 to 6M | 1.67 ± 1.45 | 1.25 ± 4.61 | 0.176 |
| | persistent CMA at 12M num inf 6 to 12M | resolved CMA at 12M num inf 6 to 12M | 1.33 ± 1.23 | 0.71 ± 0.81 | 0.108 |
| (C) | AAF num inf 0 to 6M | AAF num inf 6 to 12M | 1.56 ± 1.71 | 1.06 ± 1.18 | 0.192 |
| | AAF-syn num inf 0 to 6M | AAF-syn num inf 6 to 12M | 1.30 ± 1.52 | 0.87 ± 0.92 | 0.177 |
| (D) | AAF num inf 0 to 6M | AAF-syn num inf 0 to 6M | 1.56 ± 1.71 | 1.30 ± 1.52 | 0.688 |
| | AAF num inf 6 to 12M | AAF-syn num inf 6 to 12M | 1.06 ± 1.18 | 0.87 ± 0.92 | 0.739 |

**Figure S1.** Flowchart of the subject selection process. Red boxes: selection within the PRESTO clinical trial[3], blue boxes: selection within the Earlyfit project[2]. The final aim of the Earlyfit project is to combine data from different omics/platforms into a predictive model. Therefore samples for different omics/platforms are taken from the same subjects.

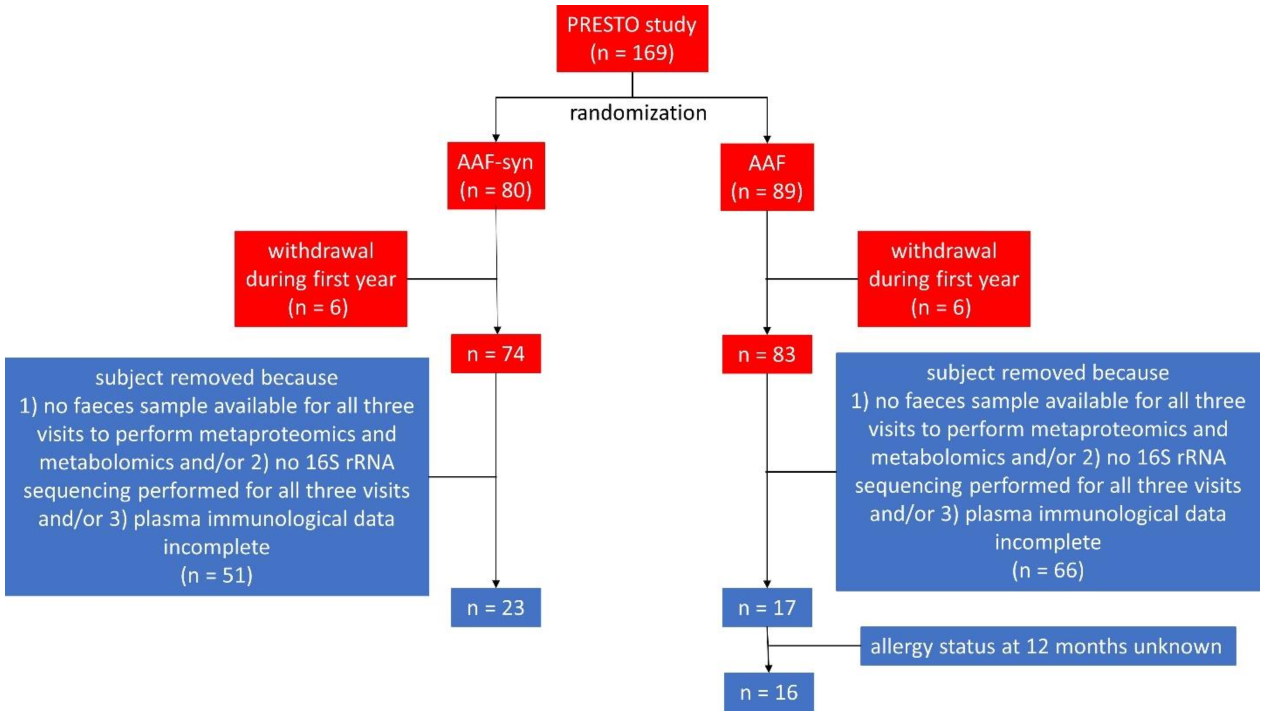


**Figure S2.** Principal components analysis colored by QC warning and showing the percentage of explained variance by PC1 and PC2.

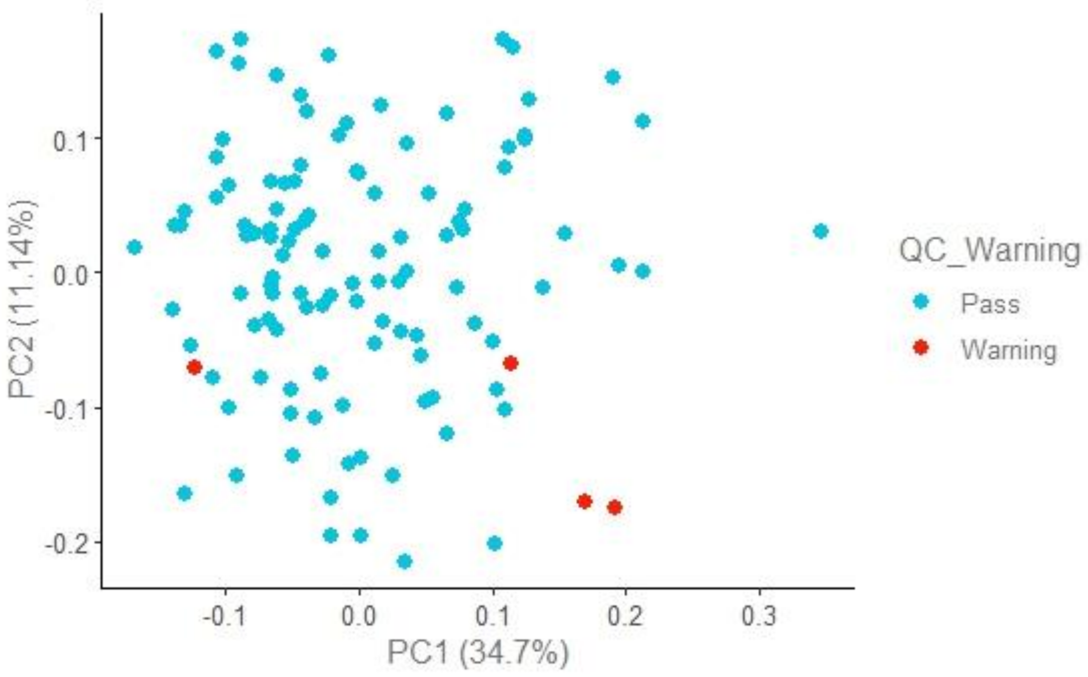

**Figure S3.** Boxplots of NPX values for the proteins with significant differences between visits within allergy groups as determined by linear mixed models and post hoc analysis. n.s: not significant (adjusted p-value > 0.05), *: 0.01 < adjusted p-value ≤ 0.05, **: 0.005 < adjusted p-value ≤ 0.01, ***: adjusted p-value ≤ 0.005.

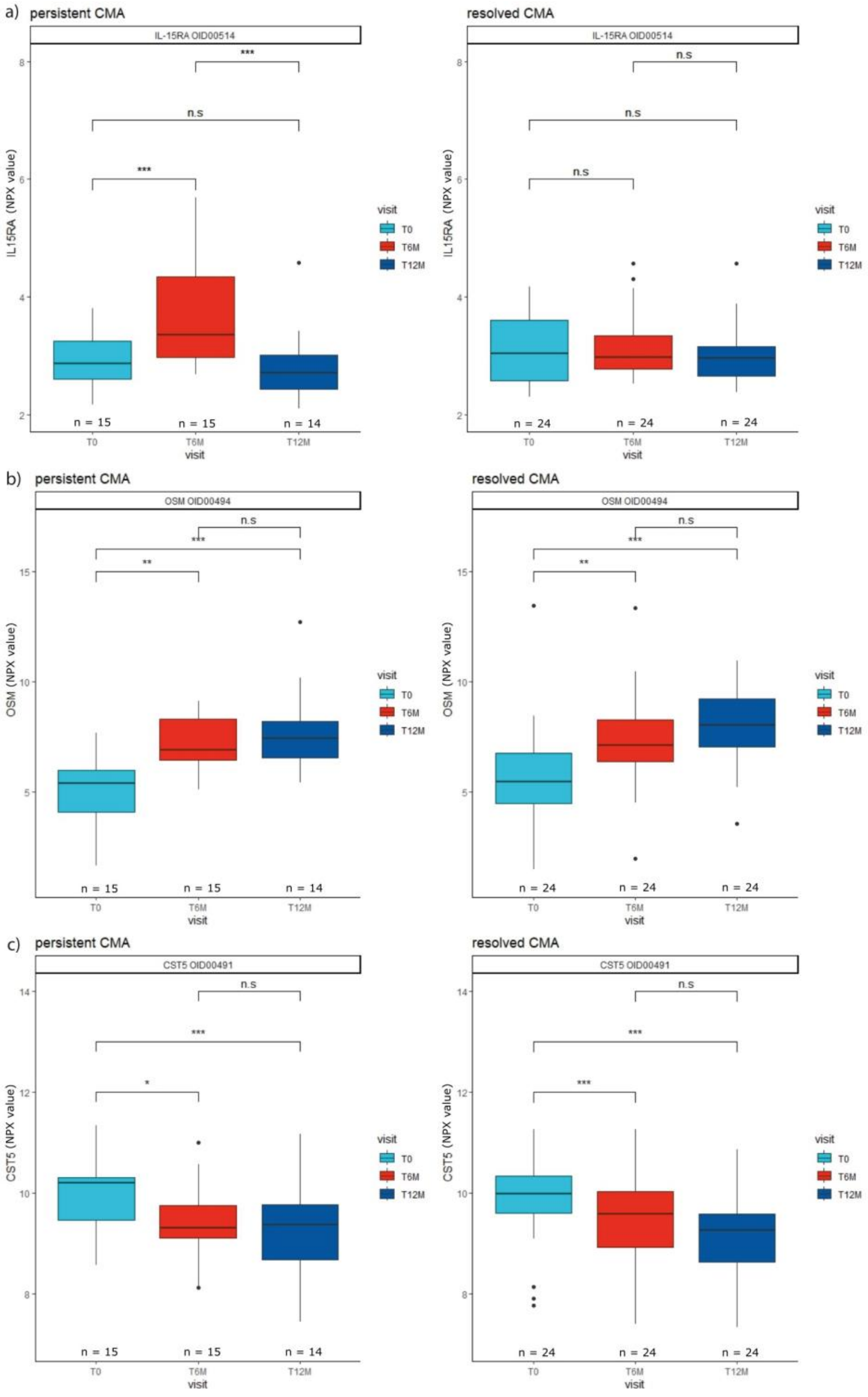

**Figure S3.** (continued)

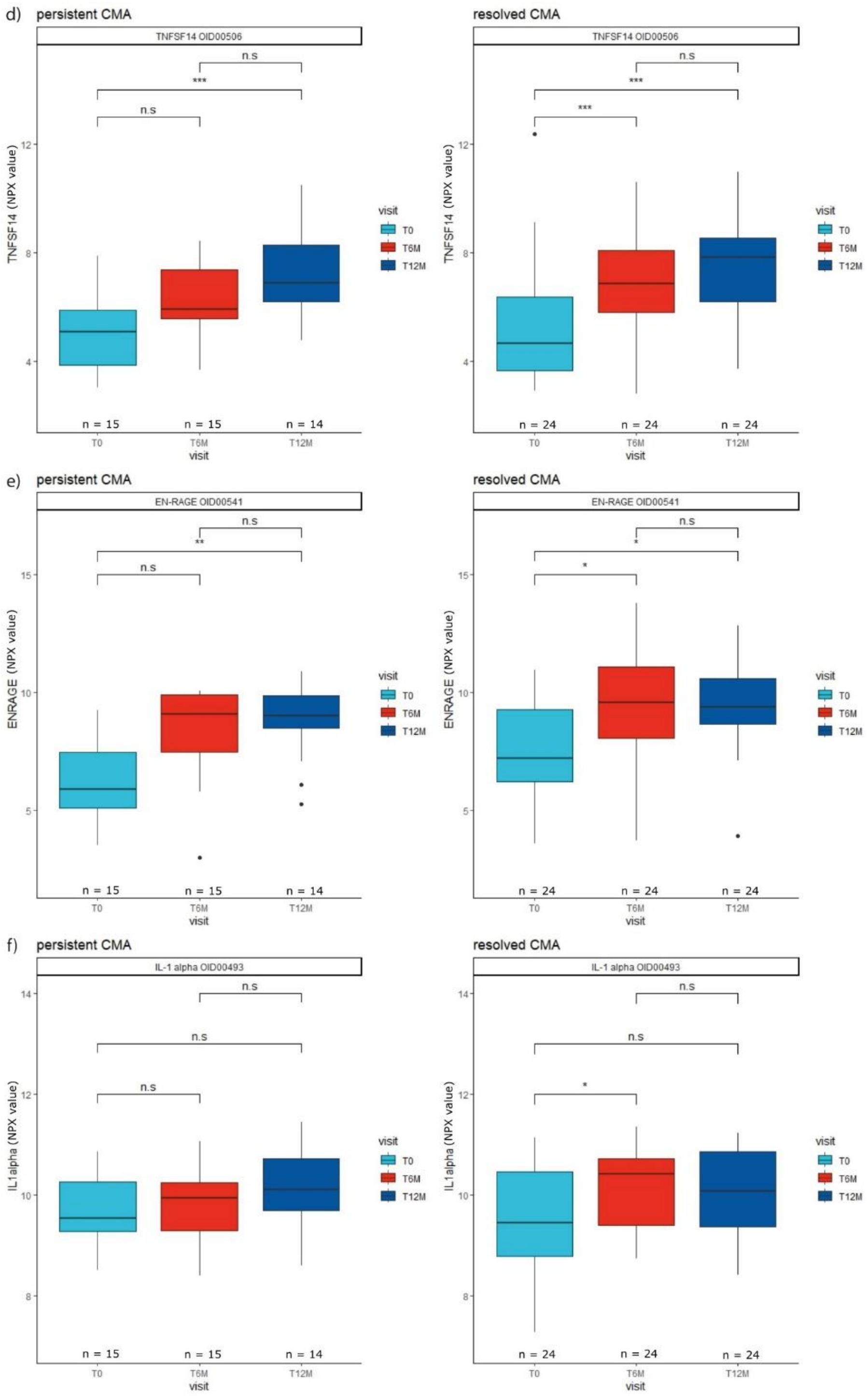

**Figure S3.** (continued)

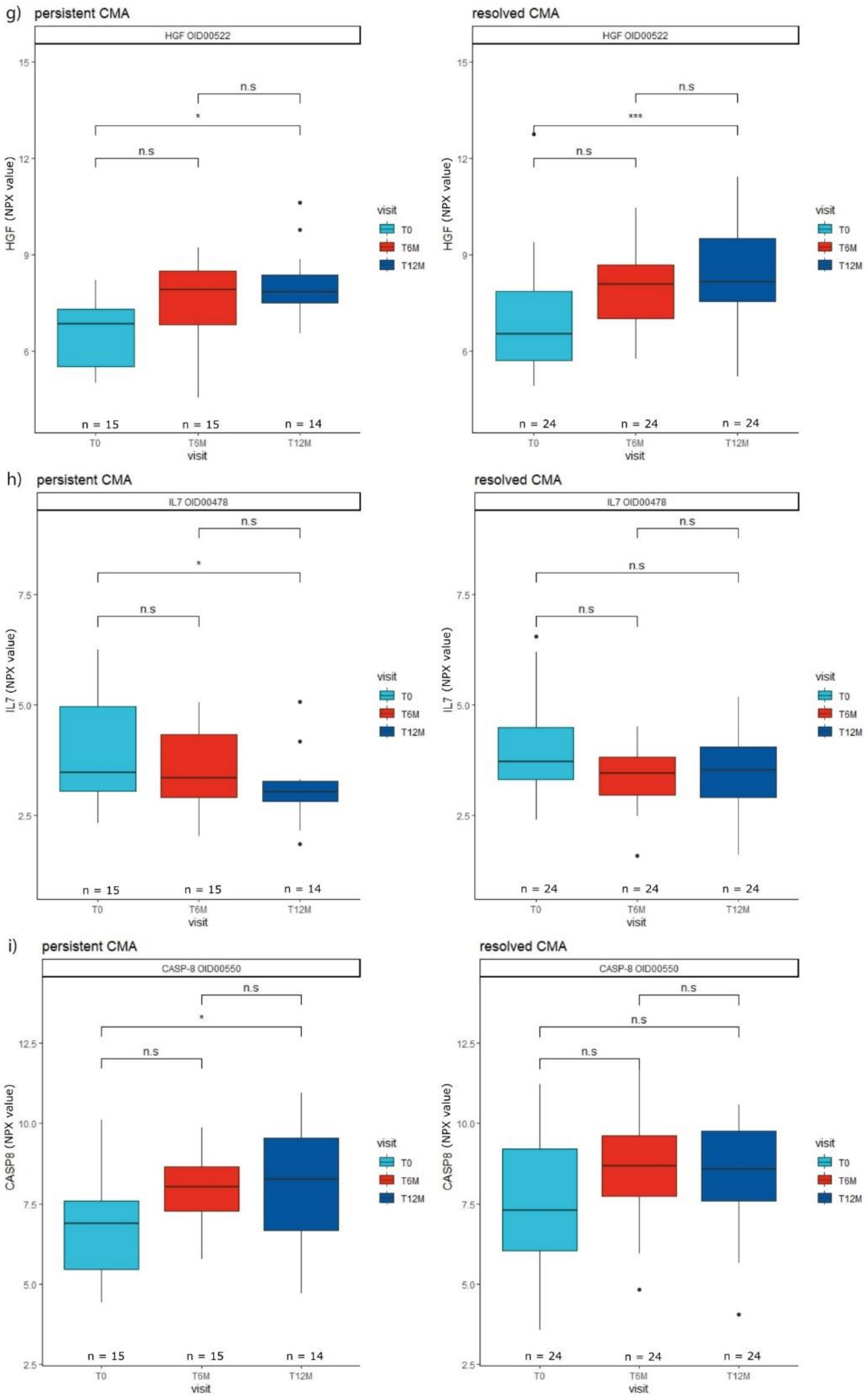

**Figure S3.** (continued)

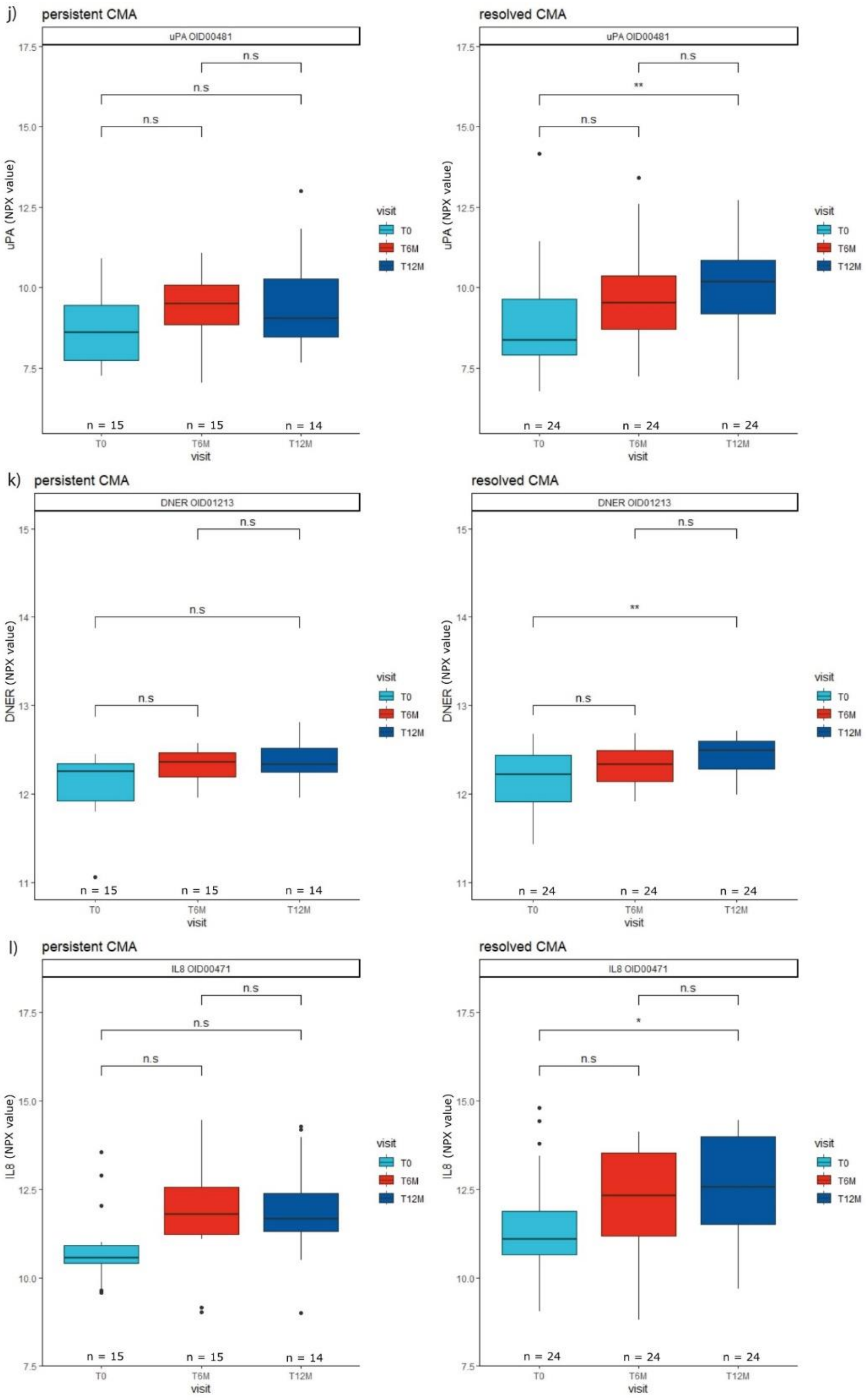

**Figure S3.** (continued)

m)

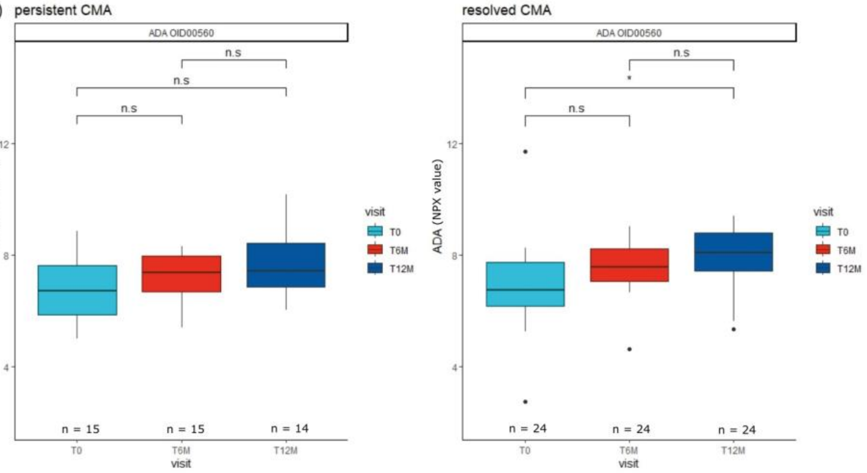

**Figure S4.** Boxplots of NPX values for the proteins with significant differences between visits within treatment groups as determined by linear mixed models and post hoc analysis. n.s: not significant (adjusted p-value > 0.05), *: 0.01 < adjusted p-value ≤ 0.05, **: 0.005 < adjusted p-value ≤ 0.01, ***: adjusted p-value ≤ 0.005.

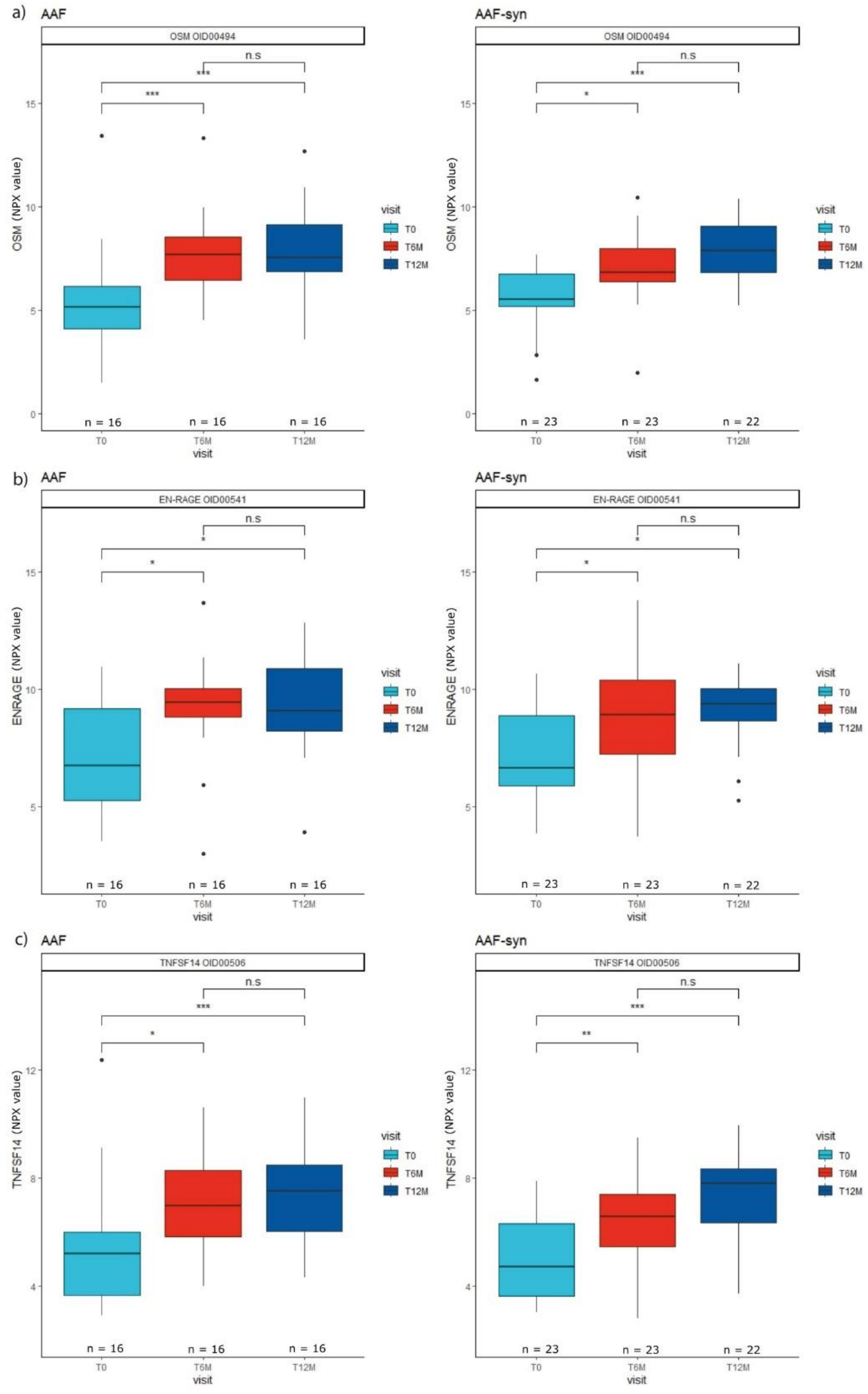

**Figure S4.** (continued)

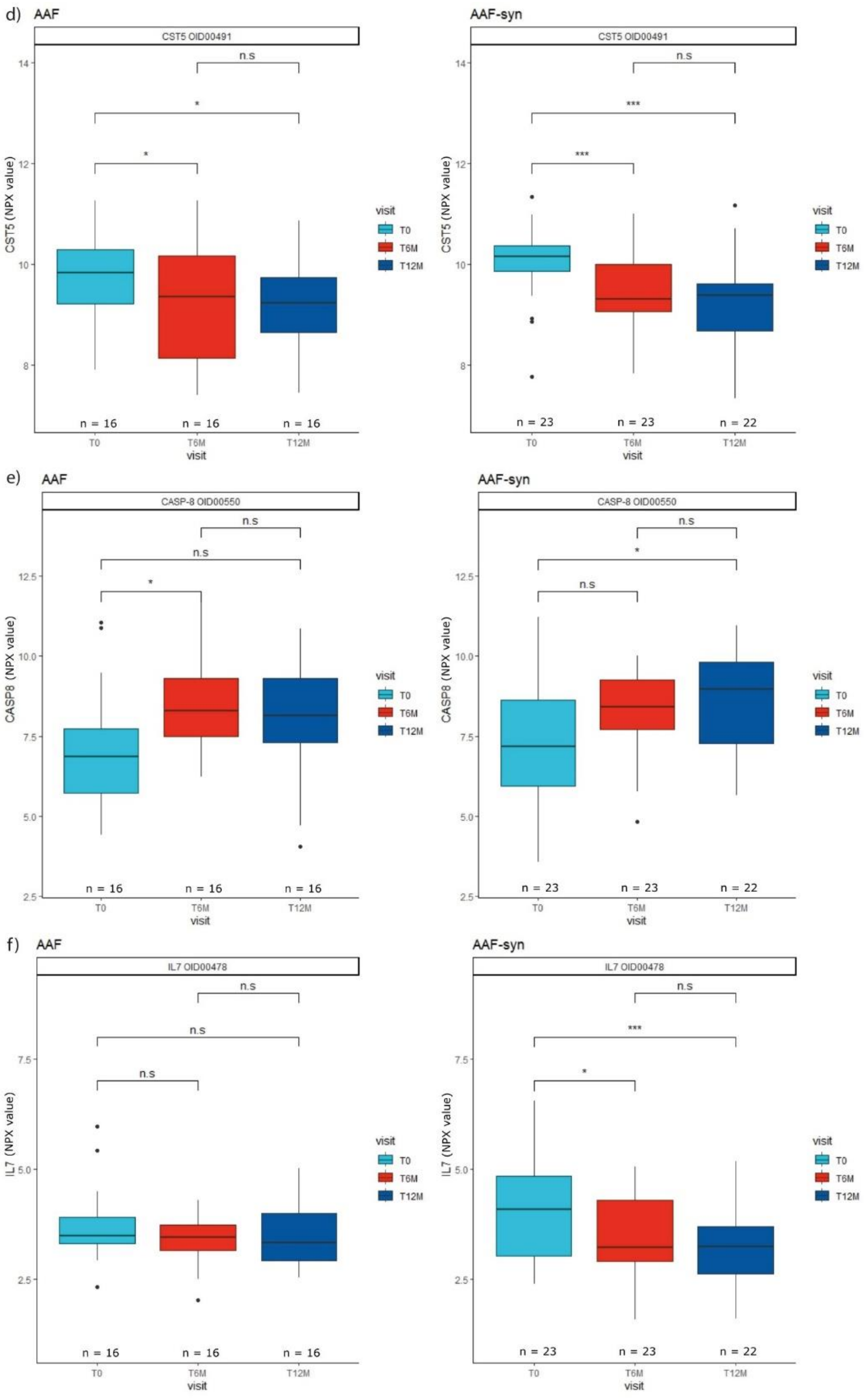

**Figure S4.** (continued)

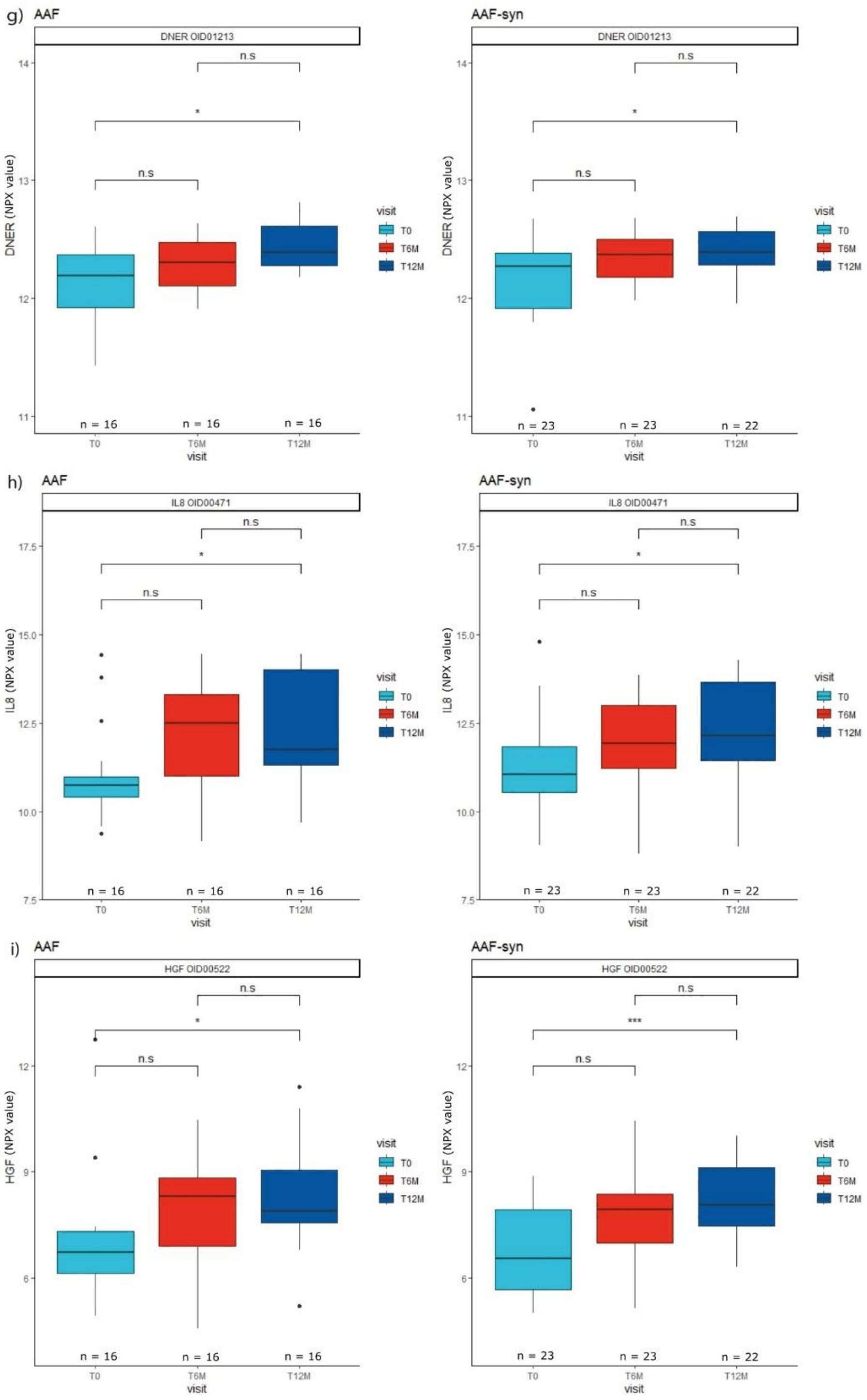

**Figure S4.** (continued)

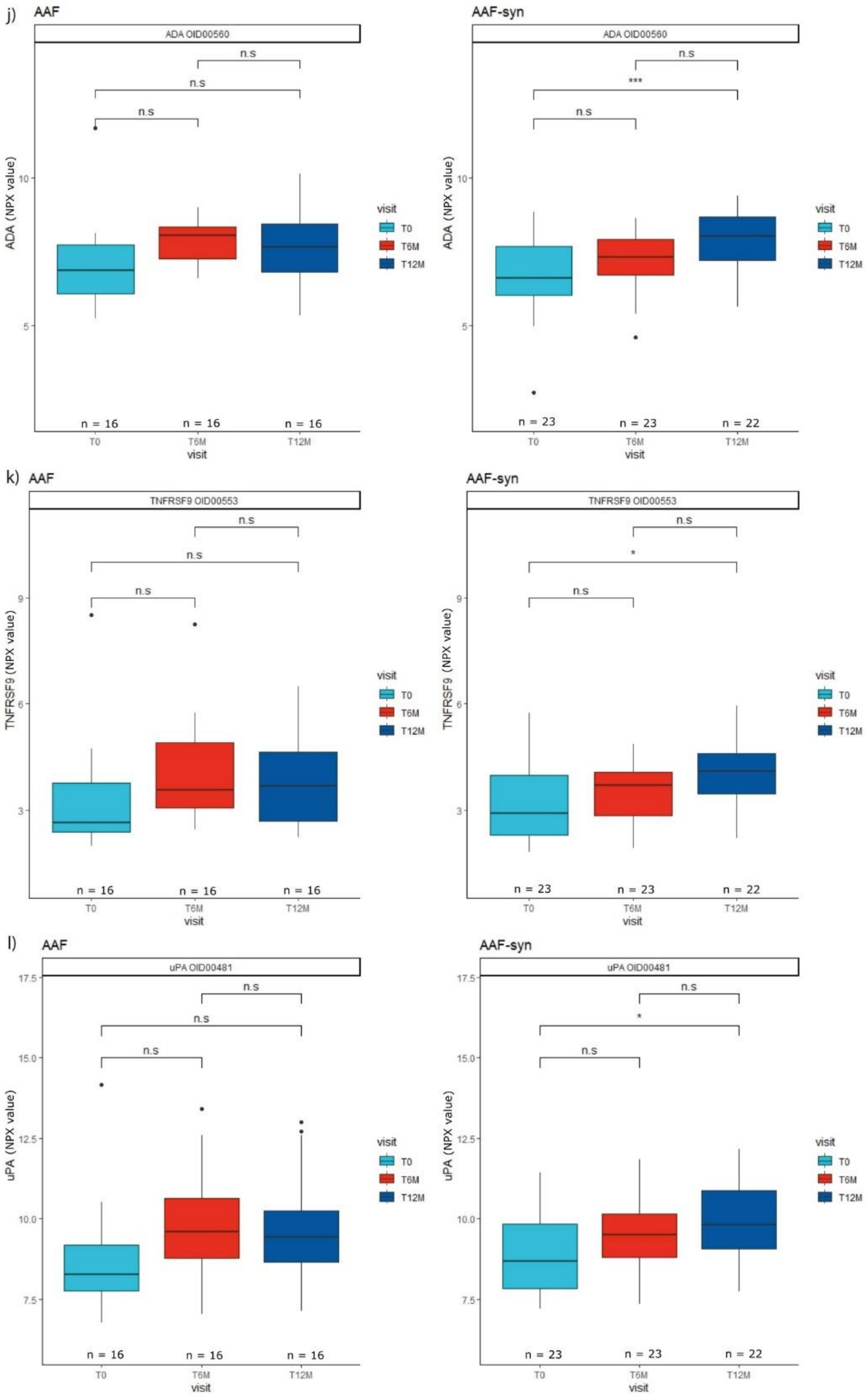

**Figure S4.** (continued)

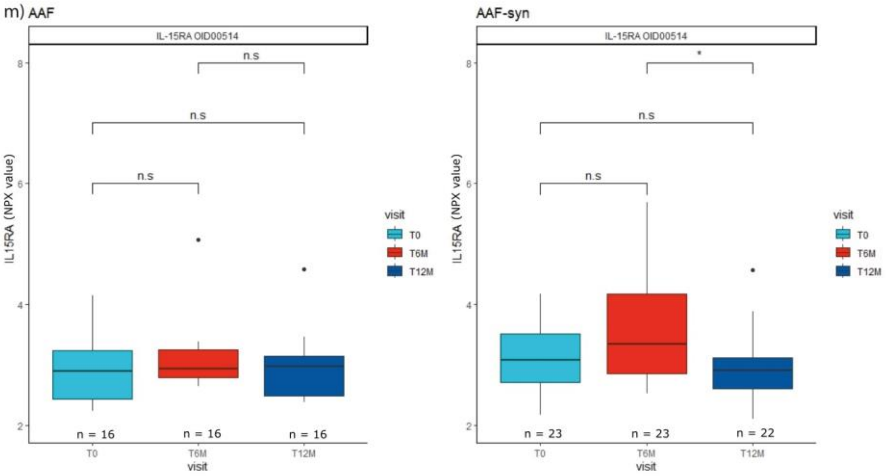


1. Benjamini Y, Hochberg Y. Controlling the False Discovery Rate: A Practical and Powerful Approach to Multiple Testing. *Journal of the Royal Statistical Society: Series B (Methodological)*. 1995;57(1):289-300. doi:10.1111/j.2517-6161.1995.tb02031.x

2. Hendrickx DM, An R, Boeren S, et al. Assessment of infant outgrowth of cow's milk allergy in relation to the faecal microbiome and metaproteome. *Sci Rep*. 2023;13(1):12029. doi:10.1038/s41598-023-39260-w

3. Chatchatee P, Nowak-Wegrzyn A, Lange L, et al. Tolerance development in cow's milk–allergic infants receiving amino acid–based formula: A randomized controlled trial. *Journal of Allergy and Clinical Immunology*. 2022;149(2):650-658.e5. doi:10.1016/j.jaci.2021.06.025
